# Studies in CRISPR-generated Mediterranean G6PD variant rats reveal G6PD orchestrates genome-wide DNA methylation and gene expression in vascular wall

**DOI:** 10.1101/2023.03.06.531429

**Authors:** Christina Signoretti, Sachin A. Gupte

## Abstract

**Background:** Recent advances have revealed the importance of epigenetic modifications to gene regulation and transcriptional activity. DNA methylation, a determinant of genetic imprinting and *de novo* silencing of genes genome-wide, is known to be controlled by DNA methyltransferases (DNMT) and demethylases (TET) under disease conditions. However, the mechanism(s)/factor(s) influencing the expression and activity of DNMTs and TETs, and thus DNA methylation, in healthy vascular tissue is incompletely understood. Based on our recent studies, we hypothesized that glucose-6-phosphate dehydrogenase (G6PD) is a modifier of DNMT and TET expression and activity and an enabler of gene expression.

**Methods:** In aorta of CRISPR-edited rats with the Mediterranean G6PD variant we determined DNA methylation by whole-genome bisulfite sequencing, gene expression by RNA sequencing, and large artery stiffness by echocardiography.

**Results:** Here, we documented higher expression of *Dnmt3a, Tet2*, and *Tet3* in aortas from Mediterranean G6PD^S188F^ variant (a loss-of-function single nucleotide polymorphism) rats than their wild-type littermates. Concomitantly, we identified 17,618 differentially methylated loci genome-wide (5,787 hypermethylated loci, including down-regulated genes encoding inflammation- and vasoconstriction-causing proteins, and 11,827 hypomethylated loci, including up-regulated genes encoding smooth muscle cell differentiation- and fatty acid metabolism-promoting proteins) in aorta from G6PD^S188F^ as compared to wild-type rats. Further, we observed less large artery (aorta) stiffness in G6PD^S188F^ as compared to wild-type rats.

**Conclusions:** These results establish a noncanonical function of the wild-type G6PD and G6PD^S188F^ variant in the regulation of DNA methylation and gene expression in healthy vascular tissue and reveals G6PD^S188F^ variant contributes to reduce large artery stiffness.

**Highlights:**

1. The wild-type G6PD and G6PD^S188F^ variant regulates the expression and activity of nuclear DNMT and TET and selectively evokes hyper/hypo-methylation of loci/promoter regions genome-wide.
2. G6PD^S188F^ variant represses and activates genes detrimental and beneficial to vascular cell phenotype and function, respectively.
3. G6PD^S188F^ variant reduces large artery stiffness.

---

Gene regulation is a formidable process now known to be mediated in part through DNA methylation and chromatin (histone) acetylation and methylation ^1^. Moreover, the impact of these epigenetic modifications on the pathobiology of cancer and vascular disease is now apparent ^2,3^. However, there are critical gaps in our knowledge regarding the epigenetic modifications [DNA methylation and histone modification] that contribute to the transformation of vascular cells from a healthy to a disease phenotype ^2^.

Several epigenetic modifiers, including histone deacetylases (HDACs), histone acetyltransferases (HATs), lysine demethylases (KDMs), and DNA methyltransferases (DNMTs) and demethylases (TETs), regulate gene expression. Interestingly, the activity of these epigenetic modifiers is driven by cellular metabolism ^4^. For example, HDAC activity depends on NAD(P)H redox ^5^, while KDM5 and TETs depend on α-ketoglutarate ^6-8^, and DNMTs depend on metabolites from ancillary one-carbon metabolism and polyamine pathways ^9^. We and others have shown that metabolites of one carbon metabolism and the polyamine pathway are significantly reduced within the lungs and smooth muscle cells (SMCs) of mice and humans after inhibition of glucose-6-phosphate dehydrogenase (G6PD) activity, as well as in red blood cells from G6PD-deficient blood donors ^10,11^. Nonetheless, the significance of G6PD in intermediary biochemical reactions, such as one carbon metabolism, to gene expression is underappreciated.

We recently discovered that G6PD interacts with epigenetic modifiers (DNMTs and KDMs) in vascular tissue ^10^, and in the lungs of hypertensive mice, inhibiting G6PD activity decreases expression of *Dnmt3b* while increasing expression of *Tet2* ^10,12^. Interestingly, induced pulmonary hypertension elicited a smaller increase in DNMT1 expression and activity in Mediterranean G6PD variant rats harboring a loss-of-function non-synonymous single nucleotide polymorphism (SNP; S188F; Type A-; G6PD^S188F^; severely deficient) than in their wild-type (WT) littermates ^13^. At the same time, DNA hypomethylation and upregulated expression of nitric oxide synthase 3 (NOS3) and superoxide dismutase were observed in the G6PD^S188F^ rats. In addition, epidemiological studies suggest that individuals in the Mediterranean region carrying a loss-of-function G6PD variant are less susceptible to cardiovascular disease ^14,15^. DNMTs and TETs affect gene expression under both normal physiological and disease conditions. Our studies indicate that levels of two metabolites in the polyamine and one-carbon pathways, spermine and dimethylglycine (DMG), respectively, are lower in vascular tissues from G6PD^S188F^ rats than their WT littermates ^16^. From those observations, we speculate that WT G6PD and the G6PD^S188F^ variant must be involved in regulating DNMT expression and activity. Up to now, however, studies have focused on how G6PD regulates the expression and activity of DNMTs and TETs under disease conditions. Its function in the normal physiological state remains unknown. Further, the extent to which G6PD controls genome-wide DNA methylation status is undetermined. In this study, therefore, we investigated the importance of G6PD to DNA methylation and gene regulation in vascular tissue from healthy WT and G6PD^S188F^ variant rats.

## Materials and Methods

### Animal models and experimental protocols

All animal experiments were approved (approval #A3362-01) by the New York Medical College Animal Care and Use Committee, and all procedures conformed to the guidelines set forth in the NIH Guide for the Care and Use of Laboratory Animals. To determine the expression and activity of DNA methylation writers and erasers in the aorta, we used loss-of-function Mediterranean G6PD^S188F^ variant rats (male: 450-950 g) and their age-matched WT littermates ^16^. Aortas were harvested and used to perform whole genome bisulphite sequencing, RNAseq, qPCR, and other biochemical analyses. Data analysis was performed in blinded fashion.

### Echocardiography

Echocardiography was performed in 2% isoflurane-anesthetized rats using a Vevo 770 imaging system (VisualSonics, Toronto, ON, Canada). Briefly, Pulse wave velocity (PWV) was determined from transit time between Doppler flow signals in the carotid and iliac arteries. Doppler signals and electrocardiogram (ECG) were recorded simultaneously, and the data were stored for subsequent off-line analysis. as described previously.^17^

### Preparation of nuclear extracts and measurement of global DNA methylation and total DNMT and TET activity

Tissue samples were pulverized in liquid nitrogen, and nuclear extracts were prepared per the manufacturer’s instructions using a kit (OP-0002-1) from EpiGentek (NY, USA). The collected nuclear extracts (20 μg protein) were then used to measure global DNA methylation and DNMT and TET activities with kits from Epigentek (USA).

### Methylated DNA immunoprecipitation (MeDIP)

In addition, we performed methylated DNA immunoprecipitation-PCR to determine overall DNA methylation status. DNA methylation was assessed using a quick EpiQuik Tissue Methylated DNA immunoprecipitation kit (Cat # P-2020, EpiGentek, NY, USA) followed by PCR according to the manufacturer’s protocol.

### Whole genome bisulphite sequencing

Genomic DNA was isolated from 5 mg of tissue, using a MasterPure™ DNA Purification Kit according to the manufacturer’s instructions. This kit employs non-enzymatically induced cell lysis followed by protein precipitation and subsequent nucleic acid isolation, which results in high yields of purified, high-molecular-weight DNA. The extracted DNA was resuspended in TE buffer and quantitated by fluorometry.

Bisulfite treatment of genomic DNA was performed using a Zymo EZ DNA Methylation Lightning Kit (www.zymoresearch.com/downloads/dl/file/id/490/d5030i.pdf). With this method, non-methylated cytosine nucleotides are converted to uracil and read as thymine (T) when sequenced. Methylated cytosines protected from conversion are still read as cytosine (C). Briefly, 50-100 ng of purified genomic DNA were treated with Zymo Lightning Conversion Reagent in a thermal cycler first for 8 min at 98°C and then for 60 min at 54°C. The bisulfite-treated DNA was purified on a spin column and used to prepare a sequencing library with an EpiGnome^TM^ Kit. In this procedure, bisulfite-treated single-stranded DNA was random-primed using a polymerase able to read uracil nucleotides to synthesize DNA strands containing a specific sequence tag at their 5′ ends. The 3′ ends of the newly synthesized DNA strands were then selectively tagged with a second specific sequence, resulting in DNA molecules di-tagged with known sequences at their 5′ and 3′ ends. Then using PCR of the original DNA strand, these tags were used to add Illumina P7 and P5 adapters at the 5′ and 3′ ends, respectively. Because only the complement to the original bisulfite-treated DNA was used as the sequencing template, the resulting Read 1 would always be the same sequence as the original bisulfite-treated strands.

The EpiGnome libraries were diluted and loaded onto a cBot DNA Cluster Generation System. Once cluster generation was complete (after ∼5 h), the flow cell is transferred to a HiSeq 2500 System for sequencing using 75-bp paired end reads. The HiSeq 2500 generates approximately 500 Gb of sequence data per flow cell or about 62 Gb per lane. Therefore, a single rat genome library can be run across two lanes of an eight-lane flow cell to generate approximately 120 Gb of data per sample. Additional sequencing can be completed for higher coverage. Methylation analysis (www.epibio.com/docs/default-source/protocols/epignome-bioinformatics-user-guide.pdf?sfvrsn=2) was performed using Bismark ^18^. FASTQ files were quality-filtered and adapter sequences trimmed using Trimmomatic ^19^. A bisulfite-converted UCSC HG19 reference genome file was generated using Bowtie 2 ^20^, after which the EpiGnome library sequence data were aligned to the reference genome. Methylation information was extracted from the output *.sam file, and genome tracks were output for visualization (performed using www.broadinstitute.org/igv/home) and reporting of downstream differential methylation calculations (performed using www.broadinstitute.org/igv/home). WGBS was performed by Genewiz.

### RNA-Seq

RNA was isolated from aorta using Qiagen® miRNeasy RNA isolation kits. In brief, 20-mg samples from each tissue specimen were flash frozen in liquid nitrogen and homogenized using a handheld homogenization pestle. Samples were then suspended in QIAzol Lysis reagent and further homogenized using a microtube homogenization mixer on ice. The subsequent steps were performed per the manufacture’s protocol. RNA samples were eluted in 50 µl of RNase free water. Adequate RNA yield was ensured using a Denovex spectrophotometer. All samples yielded greater than 350 ng/µl of RNA, after which samples were stored at −80°C until further use. RNA quality was assessed using a model 4200 TapeStation [Agilent Technologies, Santa Clara, CA], and RNA quantity was determined using a Qubit 2 fluorometer [Life Technologies, Carlsbad, CA]. For each sample, 300 ng of RNA were used to construct a cDNA sequencing library with a TruSeq Stranded Total RNA Library Preparation Kit [Illumina, San Diego, CA]. Sequencing of paired end reads [75 bp × 2] was performed in an Illumina NextSeq 550 system. Sequence reads of each sample were aligned to the reference transcriptome, and gene transcript abundance was quantified using the Kallisto [v0.43.1] algorithm [doi:10.1038/nbt.3519]. Differential expression of genes and transcripts was achieved in paired groups using the DESeq2 package [doi:10.1186/s13059-014-0550-8] on the RStudio platform [v1.3.1073, with R v4.1.0]. RNAseq was performed by Genewiz.

### Quantitative real-time PCR

Real-time PCR (RT-PCR) was used to analyze mRNA expression. Briefly, total RNA was extracted from aortas using a Qiagen miRNEasy kit (Cat # 217004). The quality and concentration of the input RNA were measured with a Synergy HT Take3 Microplate Reader (BioTek, Winooski, VT), and cDNA was prepared using SuperScript IV. VILO Master Mix (Cat # 11756500, Invitrogen) for mRNA. Quantitative PCR (qPCR) was performed in duplicate using TaqManTM Fast Advanced Master Mix (Cat # 44-445-57) for mRNA in a Mx3000p Real-Time PCR System (Stratagene, Santa Clara, CA). The primers for the qPCR were purchased from Thermo Fisher Scientific/TaqMan. mRNA levels were normalized to internal control *Tuba1a*, and relative mRNA expression was reported.

### UHPLC-MS metabolomics

Snap frozen tissues were ground to powder (GenoGrinder, SPEX, Metuchen, NJ) and extracted in ice cold methanol:acetonitrile:water (5:3:2 v/v) at a concentration of 10 mg of tissue/1 ml of solution by vortexing for 30 min at 4°C, followed by centrifugation at 15,000 g for 10 min at 4°C. Twenty microliters of supernatant were collected from each extract for metabolomics analyses. Analyses were performed using a Vanquish UHPLC system coupled online to a Q Exactive mass spectrometer (Thermo Fisher, Bremen, Germany). Samples were resolved over a Kinetex C18 column (2.1 x 150 mm, 1.7 µm; Phenomenex, Torrance, CA, USA) at 25°C using a 3-min isocratic condition of 5% acetonitrile/95% water and 0.1% formic acid flowing at 250 µl/min, or using a 9-min gradient at 400 µl/min from 5-95% B (A: water; B: acetonitrile both phases coupled with 0.1% formic acid or 10 mM ammonium acetate for positive and negative ion mode, respectively). MS analysis and data elaboration were performed as described previously ^21^. Metabolite assignment was performed using MAVEN (Princeton, NJ, USA) as described in previously ^22^.

### Measurement of G6PD activity

G6PD activity was measured spectrophotometrically using an assay purchased from Cayman Chemicals, MO. The standard assay buffer contained 1 mM MgCl_2_, 50 mM Tris, pH 8.10 (carefully adjusted with concentrated HCl), 0.1 mM NADP^+^ and 0.2 mM glucose-6-phosphate. Five-microliter aliquots (containing 5 μg of protein) of nuclear extract prepared from aorta were pipetted into the wells of a 96-well plate (Fisher Scientific) followed by addition of 200 μl of the standard assay buffer. The absorbance at 339 nm was then measured using a plate reader (Synergy HT, Biotek) immediately and for up to 25 min at 50-s intervals. Background absorbance was corrected by subtracting the value of a blank containing no homogenate from all sample readings, and G6PD activity was determined quantitatively using a molar extinction coefficient of 6220 M^-1^cm^-1^.

### Statistical analysis

N represents the number of rats per group or biological replicates. All experiments (assays) were done in duplicate (technical replicates) to ensure the reliability of single values. Statistical analyses were performed using GraphPad Prism 9 software. Values are presented as the mean ± standard error (SE). Normality and outlier identification tests were performed and outliers were removed. Statistical comparisons of samples were made for two groups with the Mann-Whitney test. To make comparisons among more than two groups, two-way ANOVA followed by Fisher’s LSD *post-hoc* test for multiple comparisons was used. Significance of RNAseq results was determined using the Benjamini-Hochberg method for multiple test correction (FDR). Values of *p<0.05* were considered significant.

## Results

### The loss-of-function G6PD^S188F^ variant led to augmented expression of genes encoding proteins involved in imprinting and negative regulation of SMC proliferation proteins and reduced expression of genes encoding proteins involved in positive regulation of cell cycle G2/M phase transition

Within cells, G6PD is localized in several subcellular compartments, including the plasma membrane, nucleus and cytoplasm ^23,24^. Within the nucleus of hepatic cells, G6PD fuels NADPH oxidase-dependent superoxide anion production ^25^, which in turn regulates gene expression in endothelial cells ^26^. To determine whether nuclear G6PD within the vasculature contributes to gene regulation, we assessed its expression and activity in nuclei isolated from rat aortas collected from G6PD^S188F^ rats and their WT littermates. We detected less (65%; p<0.05) G6PD activity in nuclei from the aortas of G6PD^S188F^ than WT aortas [Fig. 1A, B], which is consistent with earlier observations in liver tissue from G6PD^S188F^ rats ^16^ and in humans carrying the G6PD^S188F^ variant ^27^.

**Figure 1:**
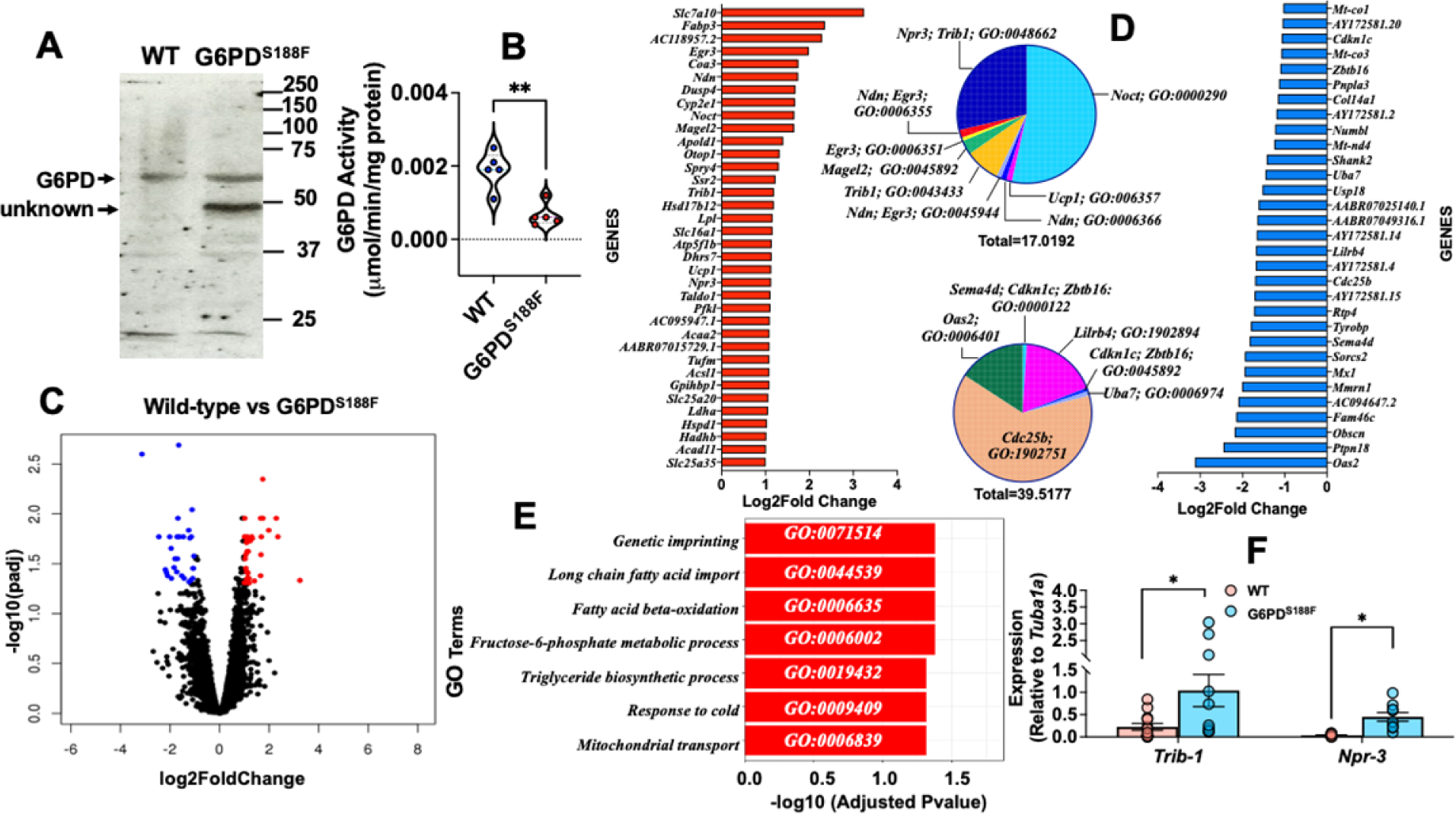
Gene expression in aortas from WT and G6PD^S188F^ rats. (A, B) Nuclear extracts prepared from aortas of G6PD^S188F^ rats show less G6PD activity but the same G6PD expression as aortic extracts from WT rats. (C) Volcano plot comparing the transcriptomic profile between WT and G6PD^S188F^ rats reveals numerous genes differentially up- or down-regulated (>1-Log2fold; black circles) between the two genotypes; for a few genes the differential increase (>1.5-Log2fold; red circles)- or decrease (>1.5-Log2fold; blue circles) was significant. (D) Top 25 increased (red bar) and decreased (blue bar) genes are shown. GO term enrichment of up-regulated (inset top) and down-regulated (inset bottom) genes are shown in a pie graph. (E) GO term enrichment of significantly altered genes is shown. (F) We confirmed the gene expression results obtained using RNAseq with qPCR and found greater expression of *Trib1* and *Npr3* in aortas from G6PD^S188F^ than WT rats. Number of rats (N) = 5 in panel A; N = 5 in panels C, D, and E; and N = 10-14 in panel F. *p<0.05 and **p<0.01.

We next performed RNAseq and found >2000 up- or down-regulated genes in aortas from G6PD^S188F^ as compared to WT rats [Fig. 1C]. However, fewer genes were significantly increased (red circle) or decreased (blue circle) over 1.5-Log2fold. GO term enrichment analysis indicated the top 25 increased genes were associated with deadenylation-dependent decapping of nuclear-transcribed mRNA, negative regulation of transcription, and SMC proliferation pathways [Fig. 1D red bars and top pie chart]. Conversely, the top 25 decreased genes were related to RNA catabolic, DNA damage, and positive regulation of cell cycle G2/M phase transition pathways [Fig. 1D blue bars and bottom pie chart]. Further, the GO term analysis also indicated significantly regulated genes were related to imprinting, long-chain fatty acid import, fatty acid β-oxidation, fructose-6-phosphate metabolic and triglyceride biosynthetic processes, response to cold, and mitochondrial import pathways [Fig. 1E]. To validate the RNAseq results, we performed qPCR for several of the genes and found that *Trib-1* and *Npr-3*, which encode proteins involved in negative regulation of SMC proliferation, were up-regulated in aortas from G6PD^S188F^ as compared to WT rats [Fig. 1F].

### The loss-of-function G6PD^S188F^ variant led to augmented expression of genes encoding DNA methyltransferases and demethylases

Because the RNAseq results revealed expression of genes encoding imprinting proteins was higher in aorta from G6PD^S188F^ than WT rats [Fig. 1] and DNA methyltransferases have been implicated in DNA imprinting ^28^, we assessed the expression and total activity of DNA methyltransferases [DNMT-1, -3A, and -3B] and demethylases [ten-eleven translocase (TET)-1, -2, and -3]. We detected 4.9-fold higher (p<0.0005) expression of *Dnmt3a* [Fig. 2B] and a concurrent 2.3-fold higher (p<0.0005) total DNMT activity [Fig. 2D] in aorta from G6PD^S188F^ than WT rats. Our results also revealed higher (p<0.05) expression of *Tet2* (2-fold) and *Tet3* (4-fold) in G6PD^S188F^ than WT aortas [Fig. 2F, G]. However, TET activity was lower in G6PD^S188F^ than WT aortas [Fig. 2H]. It is known that metabolites of ancillary one-carbon metabolism and the polyamine pathway are critical to DNA methylation and gene regulation ^9^. Because DNMT and TET activities are regulated by metabolites of the one-carbon pathway and the Krebs’s cycle [Fig. 2I], we speculated that altered activity in the one-carbon pathway and Krebs’s cycle must be contributing to the respective gain and loss of DNMT and TET activity observed in G6PD^S188F^ as compared to WT aortas. We previously showed that α-ketoglutarate – a metabolite of the Krebs’s cycle and co-factor for TET – is lower in aortas and carotid arteries from G6PD^S188F^ rats ^16^. Here, we found higher levels of methionine, a one-carbon metabolite, and higher methionine-to-dimethylglycine (DMG) ratios in aortas from G6PD^S188F^ than WT rats [Fig. 2J, K]. These results suggest the increased DNMT and decreased TET activities elicit differential DNA methylation that potentially alters gene expression in aortas of G6PD^S188F^ as compared to WT rats.

**Figure 2:**
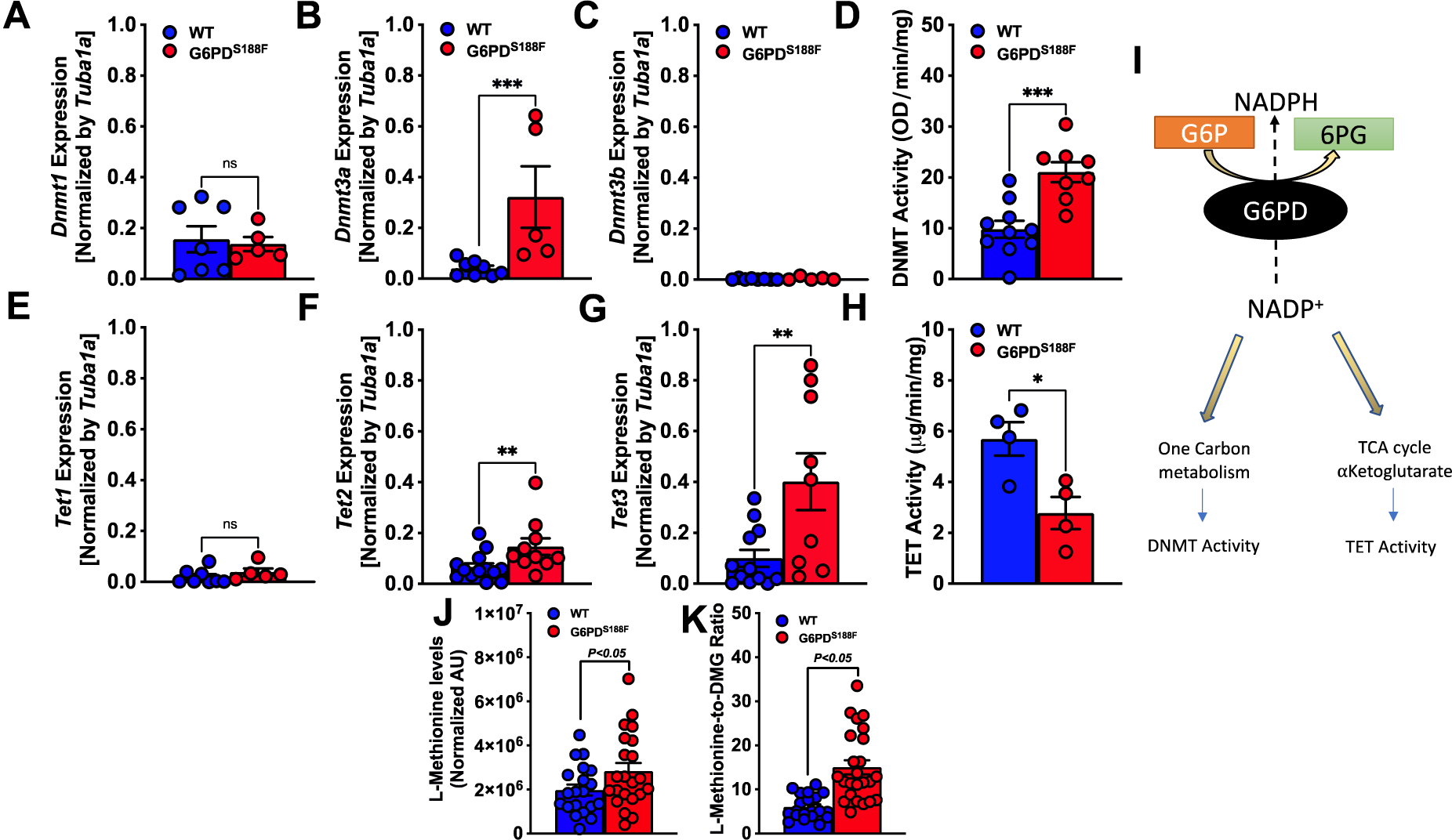
Expression and activity of DNMTs and TETs in aortas from WT and G6PD^S188F^ rats. (A to C) Determination of *Dnmt* (1, 3a, and 3b) gene expression revealed higher expression of *Dnmt3a* in aortas from G6PD^S188F^ than WT rats. (D) DNMT activity was higher in aortas from G6PD^S188F^ than WT rats. (E to G) Determination of *Tet* (1, 2, and 3) gene expression revealed higher expression of *Tet2* and *Tet3* in aortas from G6PD^S188F^ than WT rats. (H) TET activity was higher in aortas from G6PD^S188F^ than WT rats. (I) Schematic illustrating how metabolites of one-carbon metabolism and the TCA cycle regulate DNMT and TET activities. (J, K) Levels of methionine, a one-carbon metabolic intermediate that serves as a substrate for DNMT activity, and the methionine-to-dimethylglycine (DMG) ratio, are higher in aortas from G6PD^S188F^ than WT rats. *p<0.05; **p<0.005 and ***p<0.001.

### The loss-of-function G6PD^S188F^ variant led to decreases in methylated cytosine

The biological significance of DNA methylation as a major epigenetic modification affecting gene expression and cell phenotype is now well recognized. We therefore assayed global methylation levels by measuring levels of methylated cytosine (5-mC), a methylation mark, and hydroxycytosine (5-hmC), a demethylation mark, in aortic tissue from WT and G6PD^S188F^ rats. Because we found higher DNMT and lower TET activities in G6PD^S188F^ than WT aortas, we anticipated seeing higher 5-mC and lower 5-hmC levels in G6PD^S188F^ aortas. On the contrary, however, we found no change in 5-mC and higher 5-hmC content in G6PD^S188F^ than WT aortas [Fig. 3A, B]. Moreover, MeDIP assays revealed less methylation of H19-ICR gene in G6PD^S188F^ than WT aortas [Fig. 3C]. These findings indicate there may be more hypomethylated than hypermethylated loci/regions genome-wide in G6PD^S188F^ than WT aortas. Therefore, to assess the differential methylation of loci/regions genome-wide, we performed unbiased whole-genome bisulphite sequencing [WGBS] in aortas from G6PD^S188F^ and WT rats.

**Figure 3:**
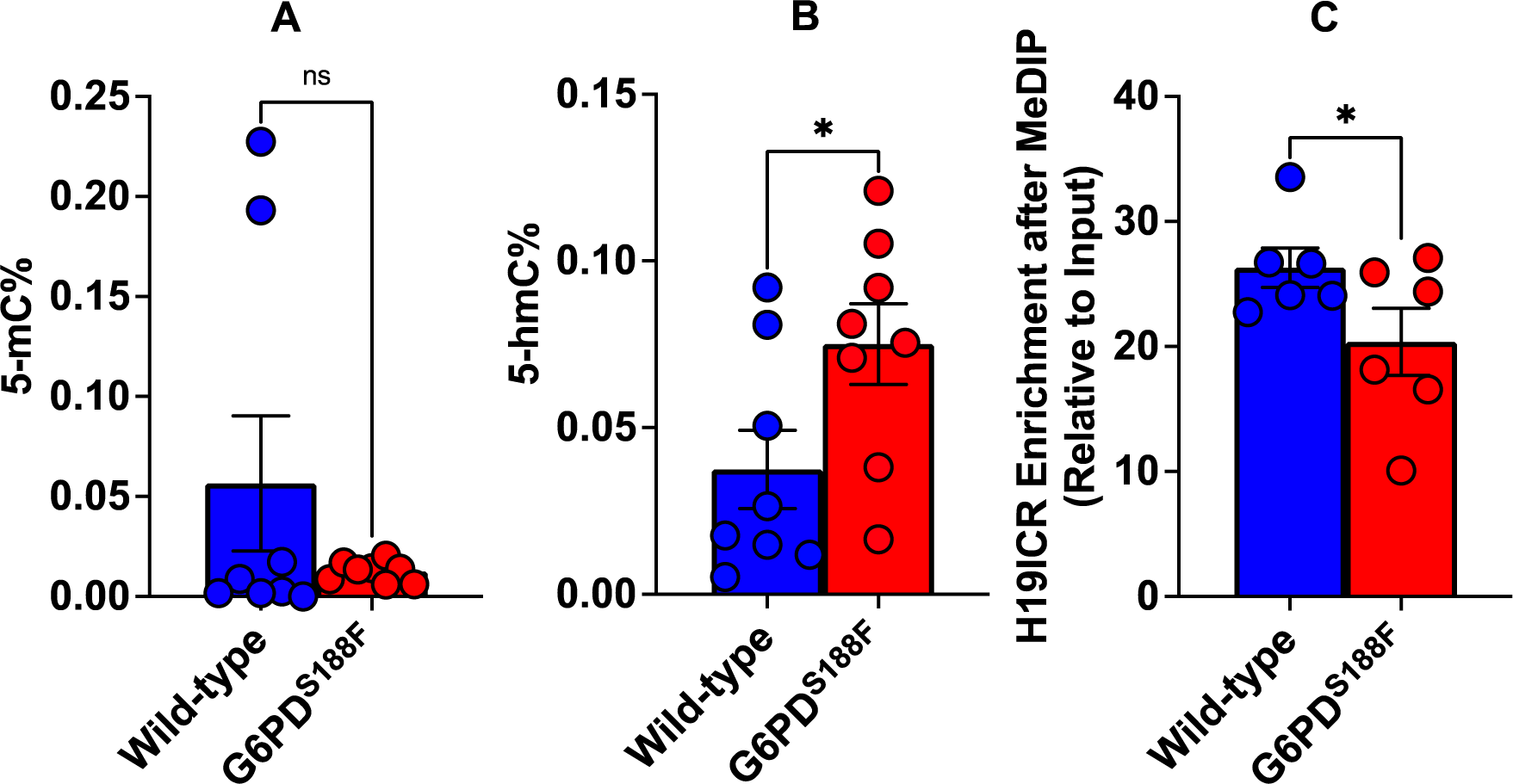
5-hmC content increased and methylation of H19-ICR gene decreased in aortas from G6PD^S188F^ rats. Determination of global methylation showed (A) 5-mC content decreased and (B) 5-hmC content increased in the aortas from G6PD^S188F^ as compared to WT rats. (C) Methylated DNA immunoprecipitation (MeDIP) assays revealed decreased methylation of the H19-ICR promoter in aortas from G6PD^S188F^ as compared to WT rats. *p<0.05 and **p<0.005.

### The loss-of-function G6PD^S188F^ variant evoked differential methylation of DNA

CpG sites with a minimum coverage of 8 reads across all samples were selected for differential methylation analysis. The summary statistics for the CpG sites’ coverage and methylation levels are given in Supplemental Figures 1 and 2. WGBS revealed 17,618 loci [of which 32% were hypermethylated and 78% were hypomethylated] and 4,222 regions [of which 42% were hypermethylated and 58% were hypomethylated] containing CpG islands that were differentially methylated between G6PD^S188F^ and WT aortas. The top 100 differentially methylated annotated loci and regions are shown in Supplemental Tables 1 and 2. The circa plot shows the start and end segments of the differentially methylated regions on each chromosome [outer track], the p-value [second track from outside], adjusted q-value [third track from outside], and % differential methylation [fourth or inner most track] in G6PD^S188F^ *versus* WT aortas [Fig. 4A]. Sequencing data then revealed that genome-wide differential methylation occurred within intergenic regions, introns, exons, and promoter regions [Fig. 4B]. Gene promoters frequently reside within CpG islands ^29^. Interestingly, among 34,525 genes examined genome-wide, we detected >25% methylation in very few promoter regions [hyper (0.037%)- and hypo (0.089%)-methylation; q-value<0.05] [Fig. 4C]. Although hyper- and hypo-methylation spanned 1,000,000 bp up- and down-stream from the transcription start site [TSS or distance from feature], most methylation changes occurred within 500,000 bp up- and downstream from the TSS [Fig. 4D]. GO term enrichment analysis indicated hypermethylated [Fig. 4E red bars] genes were related to transporter activity, molecular function regulators and transducers, protein binding activity, and catalytic activity pathways [Fig. 4F], while hypomethylated [Fig. 4E blue bars] genes encoded proteins encompassing cellular, multicellular organismal, metabolic, localization, signaling, and developmental processes [Fig. 4G].

**Figure 4:**
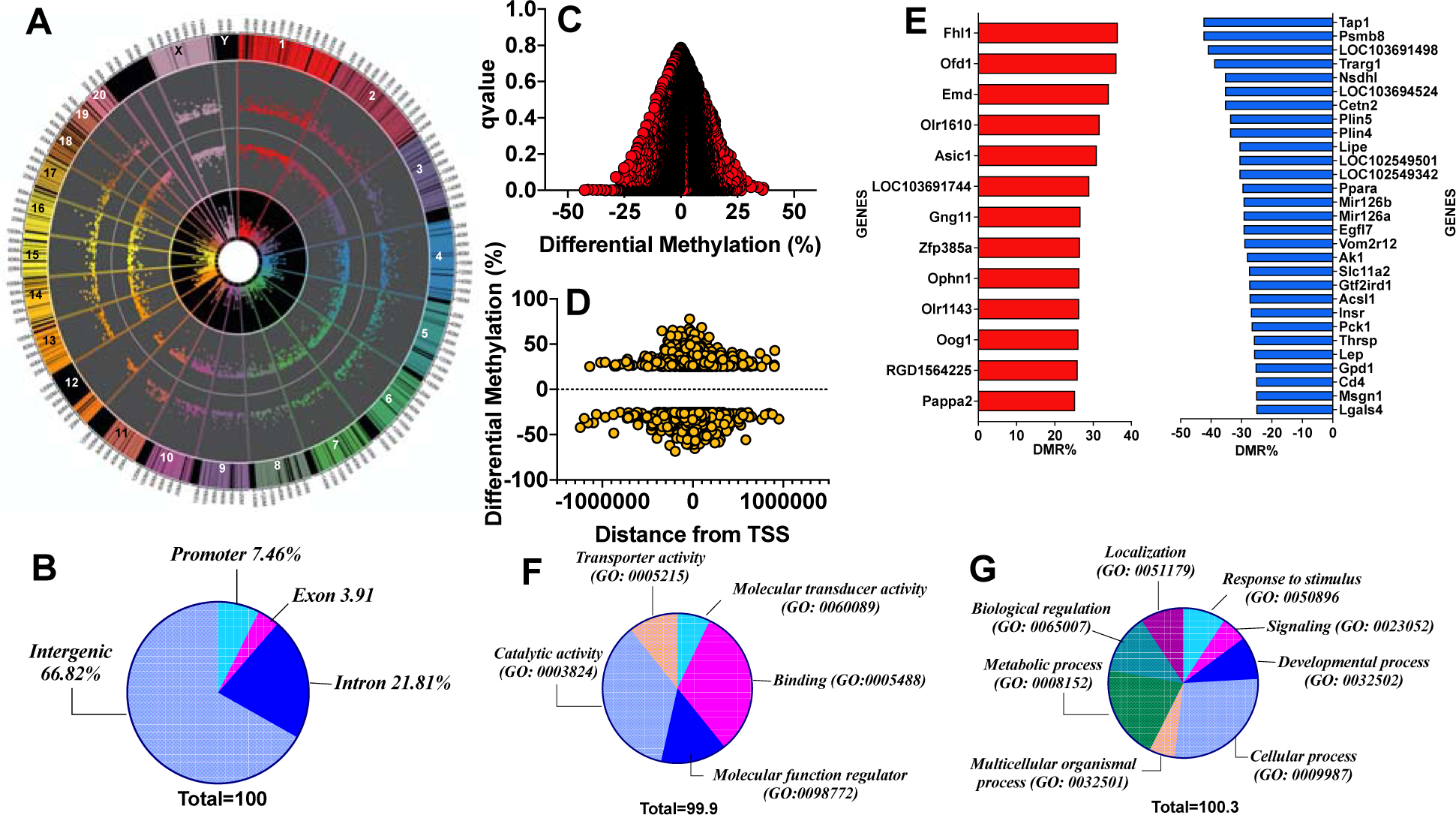
Methylome landscape in aortas from WT and G6PD^S188F^ rats. (A) WGBS in WT and G6PD^S188F^ rats revealed differential methylation of >17,000 loci genome-wide; the circa plot demonstrates differential methylation of loci on each chromosome (outer track), the padj-value (second track from outside), q-value (third track from outside), and % differential methylation (inner most track). (B) Pie chart demonstrating differential methylation of the indicated regions within the genome. Scatter plots demonstrate differential methylation of (C) promoter regions and (D) distance from the transcription start site within the genomes of WT and G6PD^S188F^ rats. (E) Significantly hyper-(red bar) and hypo (blue bar)-methylated (>25%) promoter regions are shown. Pie charts show GO term enrichment of (F) hypermethylated and (G) hypomethylated promoter regions identified in WT *versus* G6PD^S188F^ rats.

### The loss-of-function G6PD^S188F^ variant led to hypomethylated and augmented expression of genes encoding proteins involved in metabolic processes, cold thermogenesis, and reactive oxygen biosynthetic processes

Genes revealed to be differentially methylated between G6PD^S188F^ and WT aortas in the WGBS analysis were cross referenced with their gene expression profiling. The Venn diagram in Fig. 5A, shows the genes overlapping between the WGBS [differential tiles] and RNAseq data [differentially expressed genes, padj<0.05 and at least 2-fold change]. To expand the data included in the analysis, we next included all genes that were differentially expressed [padj<0.05] and were also differentially methylated in the WGBS data. That resulted in 32 genes for downstream analysis, among which expression of 29 hypomethylated genes was increased over 1-Log2fold [Fig. 5B]. GO term enrichment analysis of the genes that were up regulated and had lower methylation revealed the enriched terms to be related to fatty acid metabolism, lipid modification, lipid and organic acid catabolic processes, cold thermogenesis, ribose phosphate metabolic processes, multicellular organismal homeostasis, and reactive oxygen species biosynthetic processes [Fig. 5C, D]. The connection between the GO terms and the gene set were obtained through string analysis [Fig. 5E]. Correspondingly, fatty acid metabolites and acyl-coA products regulated by proteins encoded by *Acsl1* and *Acad11* were significantly altered in aortas from G6PD^S188F^ as compared to WT rats [Fig. 5F].

**Figure 5:**
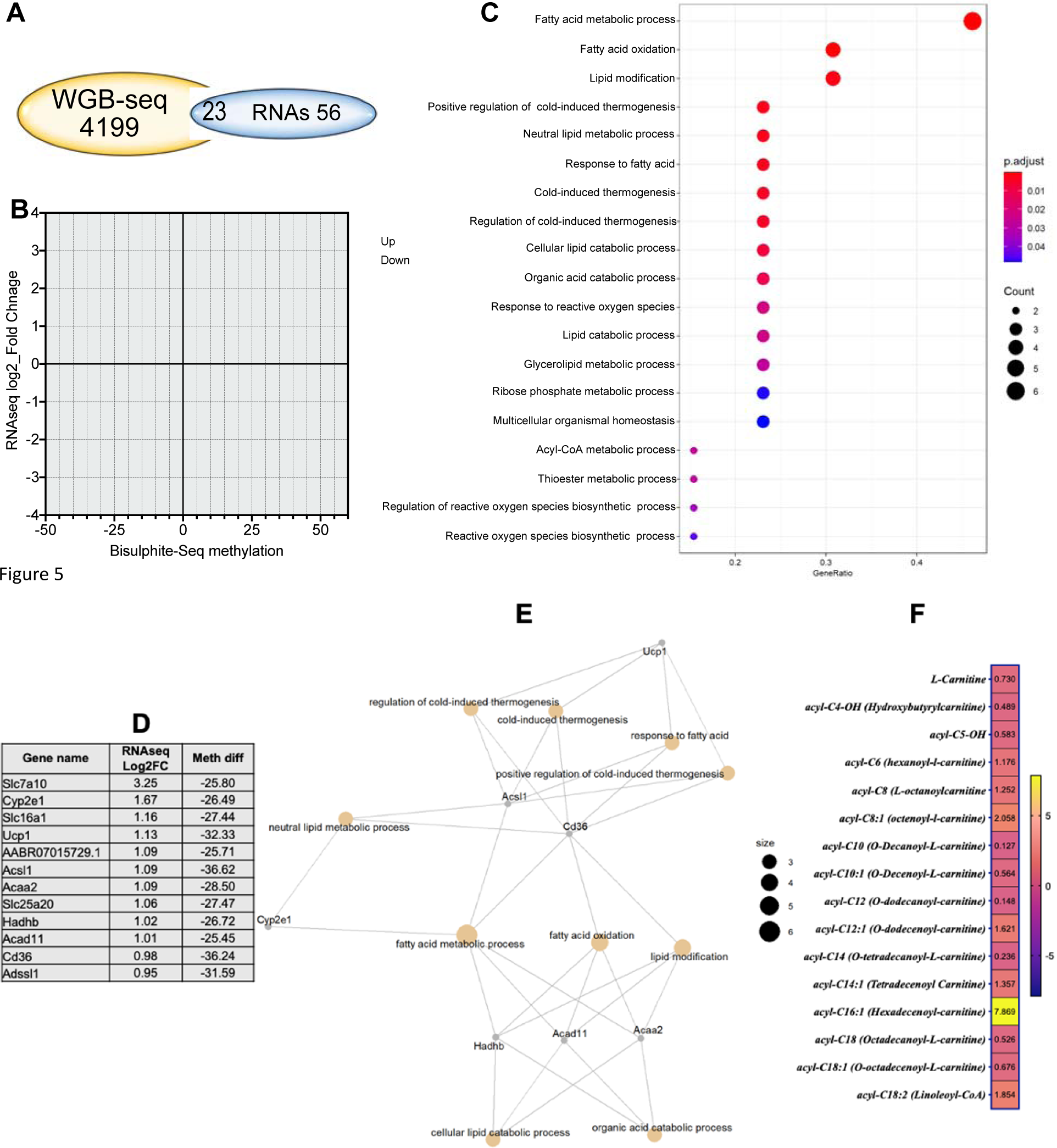
Integration of WGBS and RNAseq results reveals that genes related to metabolism and reactive oxygen species generation are affected by the G6PD^S188F^ variant. (A) Venn diagram showing the genes overlapping between the WGBS [differential tiles] and RNAseq data. (B) Thirty-two genes were differentially methylated and expressed [padj<0.05]; of those 29 were hypomethylated, and their expression was increased over 1-Log2fold. (C) GO term enrichment analysis indicating the hypomethylated and increased gene set were related to fatty metabolism, lipid modification, lipid and organic acid catabolic processes, cold thermogenesis, ribose phosphate metabolic processes, multicellular organismal homeostasis, and reactive oxygen species biosynthetic processes. (D) Percent hypomethylation and Log2fold increases in the gene set related to metabolism are shown. (E) Connection between the GO terms and the gene set were detected through string analysis. (F) Metabolomic analysis revealed fatty acid metabolites and acyl-coA products regulated by *Acsl1*- and *Acad11*-encoded proteins were significantly altered in aortas from G6PD^S188F^ rats.

### The loss-of-function G6PD^S188F^ variant led to hypomethylation of genes encoding negative regulators of SMC proliferation and steroid biosynthesis

To determine whether the G6PD^S188F^ variant leads to modification of the methylation profiles of genes involved in SMC function and/or growth, we examined the correlation between the WGBS and RNAseq data [Fig. 6A, B]. Genes related to SMC contraction [5-hydroxytryptamine receptor 1B] and SMC remodeling/inflammation [collagen alpha1 (III) and interleukin-15] were hypermethylated and down-regulated in G6PD^S188F^ as compared to WT aortas [Fig. 6C]. On the other hand, genes related to negative regulation of SMC proliferation [*Npr-3*: natriuretic peptide receptor 3] and anticoagulation [*Adtrp*: androgen-dependent TFPI-regulating protein] were hypomethylated and up-regulated in G6PD^S188F^ aortas [Fig. 6C]. qPCR performed to confirm the RNAseq results revealed higher expression of hypomethylated *Npr-3* in aorta from G6PD^S188F^ as compared with WT rats [Fig. 1F]. Further, expression of another hypomethylated gene, *Mycd*, which encodes a transcription coactivator of serum response factor that prevents dedifferentiation/proliferation of SMC, was increased in G6PD^S188F^ as compared to WT aortas [Fig. 6D]. Although our RNAseq analysis indicated expression of hypermethylated *Htr1b* and *Col3a1* was slightly decreased [Fig. 6C], qPCR analysis showed *Htr1b* expression to be increased (WT: 0.00298±0.00095 and G6PD^S188F^: 0.01259±0.00466) and *Col3a1* expression to be unchanged (WT: 35.01±6.09 and G6PD^S188F^: 42.30±9.62).

**Figure 6:**
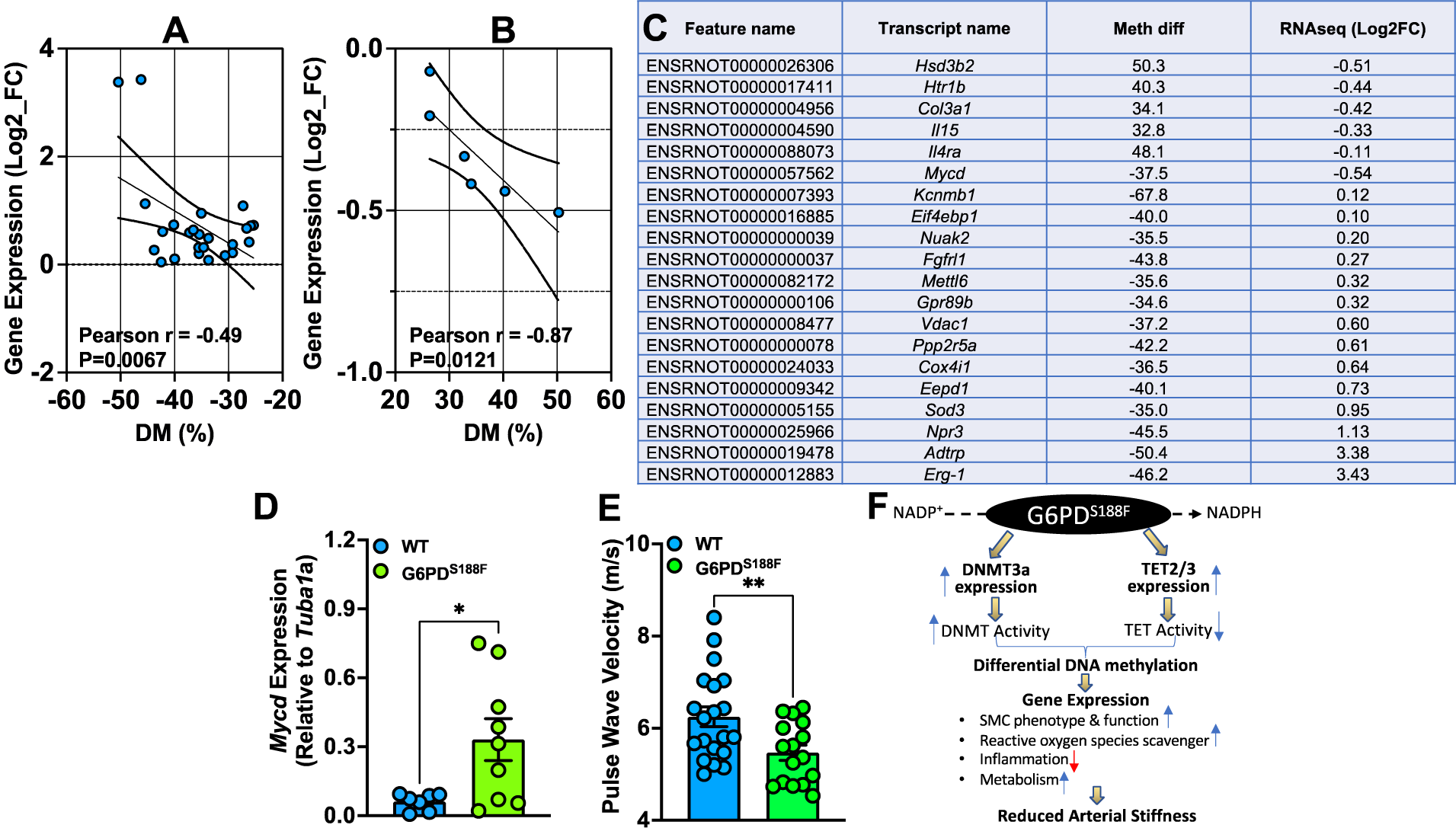
SMC phenotype- and function-related genes are differentially methylated and expressed between aortas from WT and G6PD^S188F^ rats. (A, B) In aorta, gene expression negatively correlated with % hypomethylation and hypermethylation of the loci. (C) Percent methylation and Log2fold expression of genes related to SMC phenotype and function are shown. (D) qPCR results showing increased expression of hypomethylated *Mycd* in aortas of G6PD^S188F^ as compared to WT rats. (E) Echocardiography results showing less pulse wave velocity in G6PD^S188F^ as compared to WT rats. (F) Schematic summary of the finds. *p<0.05 and **p<0.005.

### The loss-of-function G6PD^S188F^ variant reduced large artery stiffness

Next, to determine consequences of decreased inflammatory genes, and increased SMC differentiated phenotype and reactive oxygen species scavenger genes, on vascular function we measured large artery stiffness. Interestingly, G6PD^S188F^ variants as compared to WT rats had significantly less PWV [Fig. 6E].

## Discussion

This study exposed a heretofore unknown role of the G6PD^S188F^ variant connected to DNA methylation and gene regulation in vascular tissue from healthy rats. Specifically, carrying the G6PD^S188F^ variant led to hypomethylation, and up-regulation of genes associated with mitochondrial and fatty acid metabolism, cancer suppression, multicellular organismal and development processes, and SMC differentiation. Conversely, the variant led to hypermethylation, and down-regulation of genes linked to inflammation, cell proliferation, and SMC dedifferentiation. Further, our results suggest the G6PD^S188F^ variant mediated genome-wide differential methylation that activated or repressed gene transcription in all cell-types comprising vascular tissue. Notably, it activated a constellation of genes associated with differentiation of the SMC phenotype. DNA methylation, undoubtedly an important epigenetic mark of transcriptional activity, occurs in all stages of the life cycle. It is associated with the genetic imprinting [inherited DNA methylation] and gene silencing [*de novo* DNA methylation] that underlies the diverse gene expression profiles seen in the varied cells and tissues that make up multicellular organisms ^30^. The methylation of promoters residing within CpG islands results in stable silencing of gene expression during development and differentiation ^31-34^. Although DNA methylation/demethylation is dynamic and continuously ongoing in cells, the regulatory factors governing temporal control of methylation/demethylation and, in turn, gene expression are not well understood. In that regard, inhibition of G6PD activity has been shown to regulate transcription of circadian rhythm-associated genes whose expression changes in a time-dependent manner ^35^. We found that imprinted *Magel2*, which encodes a protein required for proper regulation of the circadian clock ^36,37^, was up-regulated in healthy vascular tissue from G6PD^S188F^ rats. Further, it has been proposed that a dysfunctional circadian clock contributes to the development of vascular disease ^38^. Consequently, our findings imply that G6PD is a critical contributor to the regulation of time-based DNA methylation and gene regulation, and that an imbalance in expression or activity of G6PD-dependent DNA methylation writers and erasers may contribute to the transformation of vascular cells from a healthy to a disease phenotype ^2^. In addition, these findings clarify, at least partially, why inhibiting G6PD activity or silencing *G6pd* reduces SMC dedifferentiation and the severity of some types of vascular disease, which we observed previously ^10,12,23,39-41^.

DNMT-catalyzed methylation of cytosine within CpG islands in various regions and loci within the genome regulates gene expression by recruiting proteins implicated in gene repression or by obstructing the binding of transcription factor(s) to the DNA ^30^. We recently reported that DNMT1 expression and activity are increased in the lungs of WT rats with induced pulmonary hypertension, but that effect was suppressed in G6PD^S188F^ variant rats ^13^. While *Dnmt1* was expressed 4- and 80-fold more than *Dnmt3a* and *Dnmt3b*, respectively, in vascular tissue from WT rats, *Dnmt3a* expression was higher in G6PD^S188F^ than WT rats, suggesting *de novo* hypermethylation occurring in vascular cells of G6PD^S188F^ rats facilitated the down-regulation of some, but not all, genes associated with inflammation and cell proliferation. Although there may be several reasons (e.g., increases in transcriptional activity or post-translational stability) for a selective *Dnmt3a*/DNMT3A increase in vascular cells of G6PD^S188F^ rats, our observations indicate that the increased DNMT3A potentially hypermethylated selective regions or loci within the genome of G6PD^S188F^ rats. We previously showed that G6PD and cytosine-5 methyltransferase form a complex within arterial tissue ^10^. We therefore propose that the S188F mutation, which is thermodynamically unstable and exhibits decreased thermal stability ^27^, likely alters the G6PD-DNMT3A interaction and facilitates recruitment of DNMT3A to target regions within the genome of G6PD^S188F^ rats. Studies have identified numerous DNMT-containing complexes involved in regulating DNA methylation and recruiting DNMT to the DNA ^28^. This highly diverse group of complexes plays a key role in DNA imprinting and gene silencing; for example, DNMT1-containing complexes are required for maintaining global DNA methylation following DNA replication, while DNMT3A containing complexes evoke *de novo* DNA methylation ^28^. However, the mechanism(s) by which the DNA methyltransferases and demethylases (epigenetic writers and erasers, respectively) responsible for the differential methylation of genes are recruited to specific genomic sites remains a mystery. In addition, underlying the augmented *Dnmt3a* transcription seen in G6PD^S188F^ rats was increased DNMT activity. We propose that increases in DNMT3A expression and levels of methionine, a precursor of the substrate that fuels DNMT activity, together contributed to the increase DNMT activity seen in aortas from G6PD^S188F^ rats. Collectively then, our findings indicate, for the first time, that in healthy vascular tissue, the G6PD^S188F^ variant enhances *Dnmt3a* expression and DNMT activity leading to *de novo* hypermethylation and down-regulation of some, but not all, genes detrimental to normal vascular cell physiology and function, perhaps reducing susceptibility to vascular diseases.

Active DNA demethylation of 5-mC catalyzed by TET enzymes is continuously ongoing within cells ^42,43^. Although total TET activity was decreased, our data revealed higher expression of *Tet2* and *Tet3* alongside increased 5-hmC and more hypo-than hyper-methylated loci and promoter regions in aortas from G6PD^S188F^ rats. These hypomethylated loci encompassed genes such as *Mycd* and *Kcnmb1*, which encode pro-contractile proteins ^40^; *Erg1(Sqle)*, which encodes squalene monooxygenase that regulates cholesterol biosynthesis has been associated with favorable outcome in terms of fibrous cap thickness and cholesterol efflux capacity ^44^; *Npr3*, which encodes a receptor that regulates blood pressure and inhibits SMC growth ^45,46^; and *Adtrp*, which encodes a hydrolase activity-enabling protein that prevents coronary artery disease and blood coagulation or clotting ^47,48^. We concurrently detected higher expression of those hypomethylated genes in G6PD^S188F^ than WT aortas. These unprecedented observations suggest that the elevated expression of TET2 and TET3 leads to their recruitment to selective regions of the genome in G6PD^S188F^ rats. In that regard, results from an earlier study suggest that TET2 binds to CArG-rich regions of active SMC contractile promoters (MYCD, SRF, and MYH11), and overexpression of TET2 augments the 5-hmC epigenetic landscape and contractile gene expression and attenuates intimal hyperplasia induced by wire injury to mouse femoral artery ^49^. Conversely, TET2 knockdown exacerbates vascular injury, and loss of TET2 and 5-hmC correlates with human atherosclerotic disease ^49^ and coronary allograft vasculopathy ^50^. Those studies suggest that TET2 coordinates SMC phenotypic modulation through opposing effects on chromatin accessibility at the promoters of pro-contractile-*versus* dedifferentiation-associated genes ^49^. Further, DNA methylation has been shown to prevent differentiation of non-vascular cells ^51^. Here, we demonstrate that the G6PD^S188F^ loss-of-function mutation led to augmented TET2 and TET3 expression and, in turn, hypomethylation that activated transcription of the SMC differentiation gene program as well as hypermethylation that repressed genes associated with cell inflammation and proliferation in vascular tissue. This presumably prevents transitioning of SMCs from a healthy to a disease state and reduces hypertension, large artery stiffness, and intimal hyperplasia in G6PD^S188F^ rats ^16,52^.

Although we detected higher DNMT activity and lower TET activity in aortas from G6PD^S188F^ than WT rats, analysis of global DNA methylation, MeDIP, and WGBS revealed more hypo-than hyper-methylation genome-wide in G6PD^S188F^ rats. This observation is perplexing but not unprecedented. Previous studies have shown that active DNMTs, especially DNMT3A and DNMT3B, are capable of demethylating DNA, depending on the local chromatin microenvironment ^53^. Interestingly, increasing the Ca^2+^ and H_2_O_2_ concentrations switches the enzyme activity of DNMTs from methylation to demethylation, with H_2_O_2_ inhibiting the DNA methylating activity of DNMTs by oxidizing the cysteine residues within the catalytic site ^53,54^. However, the physiological relevance of this switch remains unclear. The G6PD^S188F^ variant reduces Ca^2+^ influx into vascular cells, and likely increases oxidation conditions as a result of a decrease in the NADPH redox state and decreased clearance of H_2_O_2_ ^16^. It is also well known that G6PD-derived NADPH controls intracellular levels of H_2_O_2_ and that G6PD knockdown or inhibition increases H_2_O_2_. We therefore propose that with its increased expression, DNMT3A is recruited to selective loci/regions in the genome and that the altered redox status in the local chromatin microenvironment leads to oxidation of the DNMT3A and demethylation of the DNA. By contrast, other loci/regions in the genome likely remain methylated or hypermethylated. We further propose that the increased activities of DNMT3A, TET2 and TET3 together evoke DNA demethylation leading to more hypo-than hyper-methylation of selective loci in the genome where there is lower reducing or increased oxidative conditions in the local chromatin microenvironment, such as within the arteries of G6PD^S188F^ variant rats.

As expected, gene expression negatively correlated with methylation levels, and RNAseq and qPCR analyses revealed that expression of hypomethylated *Npr3* and *Mycd* increased. By contrast, there was a discrepancy between the RNAseq and qPCR results for hypermethylated *Htr1b* and *Col3a1*. Although RNAseq showed a small non-significant decrease in expression of hypermethylated *Htr1b* and *Col3a1*, qPCR indicated *Htr1b* and *Col3a1* expression was increased and unchanged, respectively, in aortas from G6PD^S188F^ as compared to WT rats. These results suggest that the methylation occurred in CpG regions irrelevant to transcriptional activity or, alternatively, expression of this gene may be regulated transcriptionally as well as at the post-transcriptional level.

We previously proposed that G6PD contributes to the regulation of vascular function and SMC phenotype ^39,40,55^. In those studies, we showed that increased G6PD expression and activity reduces, and G6PD inhibition or knockdown augments, expression of *Mycd* and the SMC-restricted gene program ^39^. Moreover, in endothelial cells G6PD regulates gene expression and is required for establishing vascular mural cell coverage of the dorsal aorta and blood vessel maturation during early development in both zebrafish and mice ^56^. These results underscore the significance of G6PD to the SMC phenotype, physiology, and function during development and during the post-development/adult stages of life.

The biochemical reactions and enzymes, including G6PD, that constitute to the pentose phosphate pathway are, evolutionarily speaking, very old and have likely functioned within organisms since the earliest stages of evolution ^57^. G6PD-derived NADPH, the fulcrum of redox balance, is the *sine qua non* that protects cells from oxidative stress damage and is a cofactor for numerous cellular reactions ^57^. The roles of NADPH in regulating glutathione and thioredoxin redox; NADPH oxidases, which are major sources of reactive oxygen species; thiol redox-based signaling molecules such as protein kinase G and nitric oxide synthase; and fatty acid and cholesterol synthesis are well studied ^40,57-61^. However, although the importance of subcellular G6PD and NAD(P)H redox in cardiovascular function and disease is becoming more apparent, remains poorly studied and poorly understood ^62,63^.

The results of this study have established a noncanonical function of the wild-type G6PD and G6PD^S188F^ variant in regulating the expression and activity of nuclear DNMT and TET and in evoking selective hyper/hypo-methylation of loci/promoter regions genome-wide that, respectively, repressed and activated genes detrimental and beneficial to vascular cell phenotype and function [Fig. 6F]. Consistently, we observed less large artery stiffness, which is emerging independent risk factor for cardiovascular diseases and an attractive therapeutic target ^64^, in G6PD^S188F^ rats as compared with WT rats. These findings may have significant bearing on vascular health, as the loss of G6PD function may prevent transformation of vascular cells from a healthy to a disease phenotype. As there are several G6PD SNPs in different ethnic groups world-wide, future studies in other ethnic G6PD variant models will be needed to confirm our findings in Mediterranean G6PD^S188F^ rats and determine whether the G6PD-dependent regulation of the methylome is a SNP-specific phenomenon.

## Acknowledgements

This study was supported by NLHBI (RO1HL132574 and RO1HL166546) and AHA (17GRNT33670454) to SAG.

## Author Contributions

**CS**: Performed qPCR, performed experiments, and edited the manuscript

**SAG**: Conceived the study, analyzed the results, wrote the manuscript

